# Average genome size estimation enables accurate quantification of gene family abundance and sheds light on the functional ecology of the human microbiome

**DOI:** 10.1101/009001

**Authors:** Stephen Nayfach, Katherine S. Pollard

**Affiliations:** Integrative Program in Quantitative Biology, University of California San Francisco, CA 94158; Gladstone Institutes, University of California San Francisco, CA 94158; Institute for Human Genetics, University of California San Francisco, CA 94158; Department of Epidemiology and Biostatistics, University of California San Francisco, CA 94158

**Keywords:** metagenomics, microbial ecology, human microbiome, average genome size, gut microbiome, comparative metagenomics

## Abstract

Average genome size (AGS) is an important, yet often overlooked property of microbial communities. We developed MicrobeCensus to rapidly and accurately estimate AGS from short-read metagenomics data and applied our tool to over 1,300 human microbiome samples. We found that AGS differs significantly within and between body sites and tracks with major functional and taxonomic differences. For example, in the gut, AGS ranges from 2.5 to 5.8 megabases and is positively correlated with the abundance of Bacteroides and polysaccharide metabolism. Furthermore, we found that AGS variation can bias comparative analyses, and that normalization improves detection of differentially abundant genes.

## Background

Shotgun metagenomics is increasingly being used to characterize the functional composition of microbial communities from the human body and the environment [1-4]. A common goal of these studies is to quantify gene family abundance and identify microbial genes or pathways that differ in abundance between environments, host phenotypes, or experimental conditions. Comparative metagenomics has shed light on how microbes have functionally adapted to the myriad of environments on Earth [4], and how variation in the functional composition of microbial communities can impact human health and disease [5, 6]. Functional variation can be interpreted in the context of taxonomic variation estimated from the same shotgun data, shedding light on evolutionary and ecological processes. These studies require accurate quantification of gene abundance from microbial community DNA.

While often overlooked, it is important to consider the average genome size (AGS) of microbial communities when performing comparative metagenomics analyses. From a statistical perspective, AGS can be a potential source of bias when comparing the abundance of genes between communities [7]. Specifically, the observed abundance of a gene from a community will be inversely proportional to the AGS of that community. For example, essential single-copy genes will be sequenced more often from a community with a relatively smaller AGS, because these genes make up a relatively larger fraction of the microbial genomes present in the community. If not accounted for, AGS is a source of variation between microbial communities that can generate spurious functional differences (i.e. false positives) and mask true ones (i.e. false negatives). Along with library size and gene length, AGS should be accounted for in comparative metagenomics studies.

AGS is also important for understanding the ecological and evolutionary forces acting on microorganisms within an environment. From an ecological perspective, microbial genome size may reflect environmental complexity, metabolic lifestyle, and community niche [8-12]. For example, in the gut it is believed that organisms with larger genomes follow more generalist lifestyles whereas those with smaller genomes are more specialized [8]. To illustrate this, *Bacteroides thetaiotamicron* (6.5 Mb) has a large genome with the metabolic potential to utilize a variety of both diet and host-derived glycans, whereas *Methanobrevibacter smithii* (1.9 Mb) is a methanogen specialized for utilization of H_2_. From an evolutionary perspective, genome size may reflect the degree of genetic drift, population size, and horizontal gene transfer within a community. For example it is believed that smaller population sizes and greater genetic drift can lead to fixation of deletions and decreases in genome size [13], whereas larger population sizes weed out these mutations and can promote gene-gain via horizontal gene transfer.

However, up to this point it has not been possible to rapidly and accurately estimate the AGS of microbial communities due to a lack of software designed for short-read metagenomics data. This has limited our understanding of both the extent and impact of AGS variation in many environments including the human microbiome. The only available software tool, GAAS [10], estimates AGS based on BLAST searches of shotgun sequences against a database of microbial genomes. Given the large and increasing scale of both metagenomics data and reference genomes, and the relatively slow speed of BLAST, this method is not computationally practical. Additionally, microbial communities frequently contain high fractions of “novel” organisms, which have not been cultured or sequenced. Even in the well-studied gut microbiome, it has been estimated that on average 43% of species abundance and 58% of richness cannot be captured by current microbial reference genomes [14]. It is not clear whether GAAS is able to accurately estimate AGS for metagenomes composed of novel taxa. Raes et al. [9] proposed to address these issues by estimating genome size based on the density of reads assigned to essential single-copy genes using BLASTX. While significantly faster and less dependent upon community composition, this method was not designed for reads shorter than 300 base pairs (bp) and no software was released. Others [15, 16] have described similar methods, but these have not been extensively validated or made available as software.

To address this problem, we developed MicrobeCensus to rapidly and accurately estimate AGS from shotgun metagenomics data and applied our tool to over 1,300 human microbiome samples. We adopt a similar approach to Raes et al., but make significant improvements which allow us to rapidly estimate AGS using reads as short as 50 bp. We find that AGS differs significantly both within and between body sites in the human microbiome and tracks with major functional and taxonomic differences between communities. For example, in the gut, AGS ranges from 2.5 to 5.8 megabases (Mb) and is positively correlated with the abundance of *Bacteroides* and genes related to polysaccharide utilization. Furthermore, we confirmed that AGS is a major bias in comparative metagenomic analyses, and that normalization improves detection of differentially abundant genes.

## Results and discussion

### Estimation of average genome size from metagenomic data

At the core of our method is a database of 30 essential single-copy gene families found in nearly all Bacteria and Archaea (Additional file 1) [17]. Because of this property, the number of shotgun sequences from these genes is expected to be directly proportional to the total coverage of microbial genomes present in a metagenome. Likewise, the fraction of total shotgun sequences that are from these genes is expected to be indirectly proportional to the average size of genomes present in a metagenome. For example, these essential genes will be sequenced more often from a community with a relatively smaller AGS, because these genes make up a relatively larger fraction of the microbial genomes present in the community.

Our method leverages this simple relationship to estimate AGS from metagenomic data (Figure 1 and Methods). In brief, MicrobeCensus takes a sample of *n* reads of length *i* from a metagenomic dataset and performs a translated alignment of these reads against the database of essential genes. Reads are classified into an essential gene if their top-scoring alignment meets or exceeds a set of minimum criteria (bit-score, alignment coverage, percent identity), which we rigorously optimized for each gene across a range of read lengths (50–500 bp). This optimization significantly improves performance by limiting the technical variation of these genes across metagenomes, and is particularly important for read lengths shorter than 100 bp (see methods). Based on the fraction of reads assigned to each gene, we then estimate AGS using one-parameter linear models, which have been trained from extensive metagenomic simulations. The purpose of this model training is to quantify for each essential gene the constant of proportionality between the number of reads assigned to that gene and the total coverage of genomes in the sample at a giving read length *i* and library size *n*. Finally, we remove any outlier predictions, and take a weighted average over the remaining estimates to produce a robust estimate of AGS.

**Figure 1.**
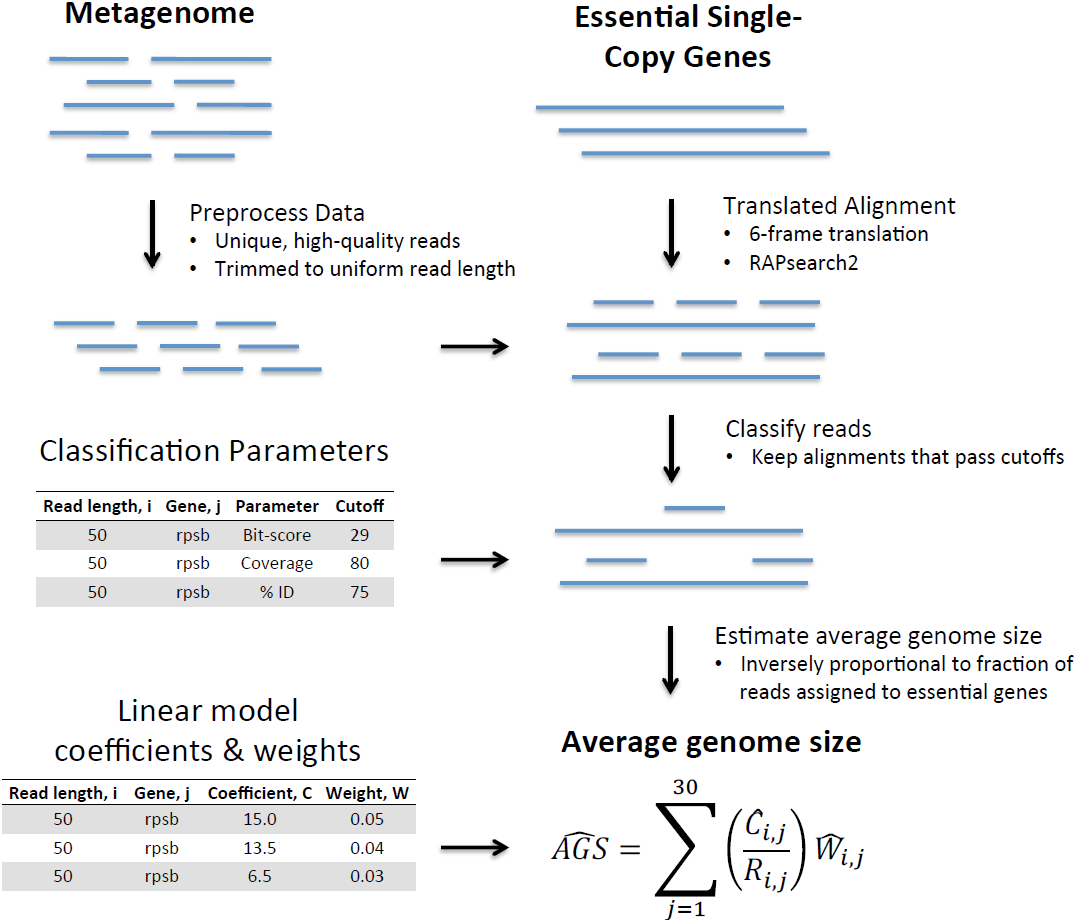
Estimation of average genome size. To estimate average genome size, MicrobeCensus takes a sample of *n* unique, high quality reads from the input metagenome and trims them to a uniform read length, *i* bp. Next, MicrobeCensus performs a translated search of these reads against the database of 30 essential genes. A read is classified into essential gene family *j* if its top-scoring alignment meets or exceeds a set of minimum criteria (bit-score, alignment coverage, percent identity), optimized for gene *j* and read length *i*. Based on the fraction of reads assigned to each gene, *R*_*i,j*_, we then estimate average genome size (AGS) using simple, one-parameter linear models, in which AGS is indirectly proportional to *R*_*i,j*_, by way of a constant of proportionality, *C*_*i,j*_. Finally, MicrobeCensus removes any outlier predictions, and takes a weighted average over the remaining estimates to produce a robust estimate of AGS.

MicrobeCensus is provided as a command line software package written in Python. Software, examples and documentation are freely available at: https://github.com/snayfach/MicrobeCensus. All dependencies are included in the software distribution.

### Comparison to existing methods

Because MicrobeCensus estimates genome size using a set of genes present in nearly all Bacteria and Archaea, we hypothesized that our tool would be robust to high proportions of novel taxa present in metagenomes. Conversely, we suspected methods that rely on reference genomes to estimate AGS would not perform as well in these cases.

To this end, we benchmarked MicrobeCensus against the tool GAAS for 20 randomly selected simulated shotgun libraries (Additional file 3). Each library was composed of 150-bp reads from a single organism. We estimated AGS at different levels of “taxonomic exclusion” in order to simulate libraries composed of organisms from novel taxonomic groups. This meant that we discarded alignments where the target reference genome or gene originated from the same taxonomic group (i.e. species, genus, etc.) as the organism in the simulated dataset. Additionally, for MicrobeCensus, we retrained our linear models while excluding all training data that originated from the same taxonomic group as the organism in the simulated dataset.

Without any taxonomic exclusion (i.e. allowing sequence data from test genome to be present in the reference database), both GAAS and MicrobeCensus produced very estimates of genome size for the 20 datasets (Figure 2a). However, when the test genome was excluded from the reference database (i.e. strain or species level taxonomic exclusion), the median unsigned error for GAAS increased to 9.5%, while the median prediction error for MicrobeCensus remained below 2%.

**Figure 2.**
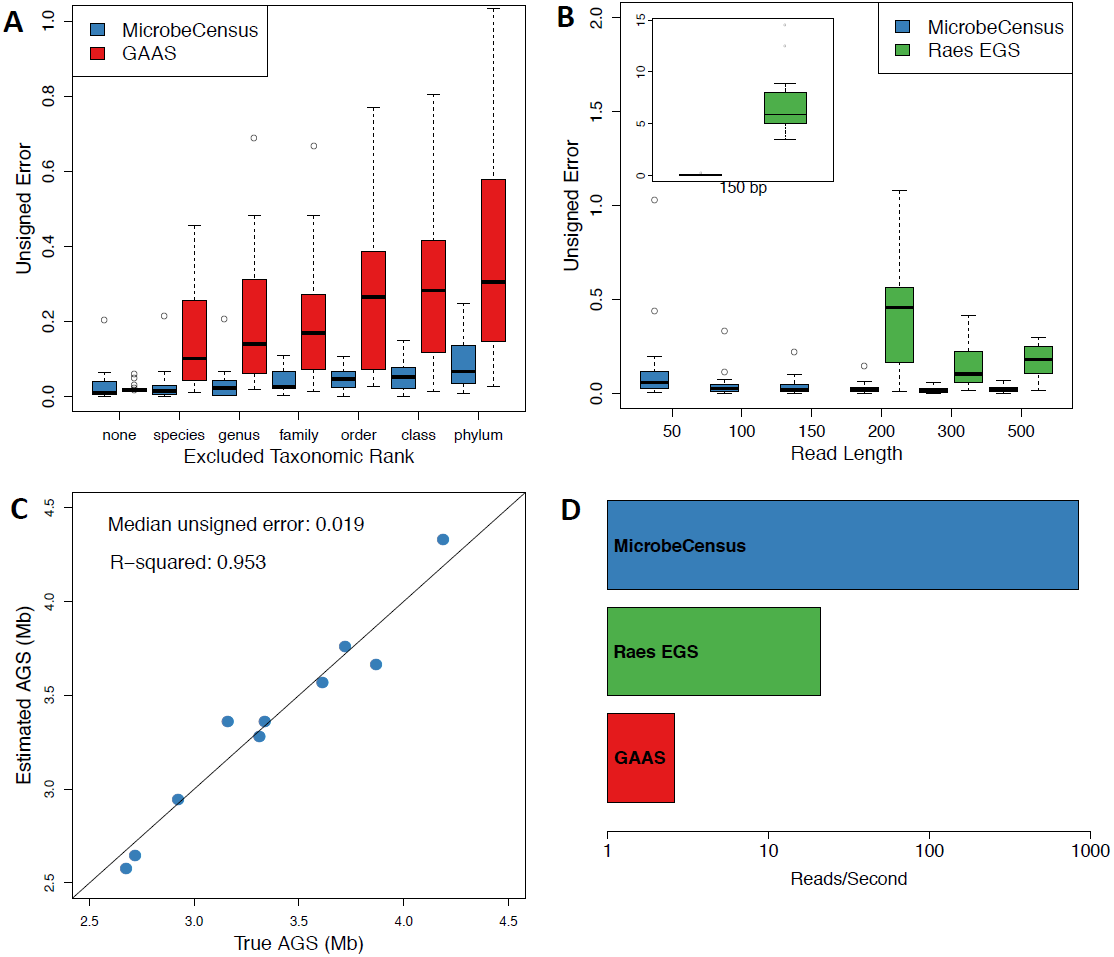
Prediction accuracy of MicrobeCensus. Performance and speed of MicrobeCensus were compared to existing methods for real and simulated data. For all comparisons, unsigned prediction error is defined as |estimated AGS – true AGS|/true AGS. **(a)** Prediction error for MicrobeCensus versus GAAS for 20 simulated shotgun libraries across different levels of taxonomic exclusion. **(b)** Prediction error for MicrobeCensus versus the method described by Raes et al. for 20 simulated shotgun libraries across read length. No estimates of genome size could be obtained by the Raes method for libraries ≤ 100 bp. **(c)** Prediction accuracy for MicrobeCensus on 10 metagenomes composed of real Illumina reads. **(d)** Speed (reads/second) of MicrobeCensus compared to GAAS and the method described by Raes et al. on a simulated 150-bp library.

Confirming our hypothesis, MicrobeCensus was able to generate reasonable estimates of genome size even for organisms from completely novel phyla (6.7% median unsigned error), while the median prediction error for GAAS was over 30%.

Additionally, we compared the performance of MicrobeCensus against the method described by Raes et al. [9], who were the first to use essential single-copy genes to estimate genome size from metagenomic data. However, this method only considered one set of alignment criteria (≥60 bits and ≤50 percent identity) for determining if a metagenomic read originated from an essential gene. While the authors were able to demonstrate accurate prediction of genome size from metagenomic data, performance was not evaluated for reads shorter than 300 bp. We hypothesized that this method would not be able to accurately estimate genome size for modern short-read libraries.

To evaluate this, we implemented the method described in Raes et al. and benchmarked this method against MicrobeCensus using simulated shotgun libraries composed of reads ranging from 50 to 500 bp (Figure 2b). We found that the Raes method was unable to generate any estimates of genome size for the short-read libraries (≤ 100 bp) due to a complete lack of alignments that exceeded the method’s classification criteria. In other words, these parameters were far too strict for the short-read libraries. For intermediate read lengths (150–200 bp), the method was able to generate estimates of genome size, but was not accurate (46–587% median unsigned error) and tended to overestimate size. As expected, long-read libraries (300 - 500 bp) produced moderately accurate genome size estimates (10-18% median unsigned error), although tended to underestimate size. In contrast, MicrobeCensus was able to accurately estimate genome size across all libraries, using reads as short as 50 bp (all < 6% median unsigned error).

### MicrobeCensus is robust to technical errors in real shotgun libraries

While we were able to obtain accurate estimates of genome size for simulated libraries, it was not clear if we could accurately predict AGS from real libraries that contained sequencing error, duplicate reads, and a non-uniform distribution of coverage. Additionally, we wanted to estimate prediction error when libraries were composed of many organisms.

To evaluate this, we identified and downloaded 42 Illumina WGS datasets from completed microbial sequencing projects where the sequenced organism was not used to train our method (Additional file 4). After removing adaptor contamination, duplicate reads, and extremely low quality reads, we pooled this data to create ten mock metagenomes (Additional file 5). Each metagenome was composed of 5 million, 70-bp reads from ten randomly selected genome projects (i.e. 500,000 reads from each project). The AGS of these mock metagenomes ranged from 2.6 to 4.1 Mb. We used MicrobeCensus to estimate AGS for each of these ten mock metagenomes (Figure 2c). Overall, we estimated AGS with 1.9% median unsigned error and less than 0.1% median signed error, indicating that we are able to obtain accurate, unbiased estimates of AGS from real metagenomic data. We also used MicrobeCensus to estimate AGS for each of the individual genome projects. However, here, our prediction accuracy was not quite as good (6.4% median unsigned error, −0.5% median signed error), suggesting that it is actually easier to predict AGS from datasets with multiple organisms (Additional file 4).

Finally, we evaluated the effect of various quality control procedures on AGS prediction accuracy. Interestingly, we found that while removing adaptor contamination and filtering duplicate reads resulted in a marginal reduction in prediction error, quality-filtering reads had very little benefit and in some cases actually reduced accuracy (Additional file 6). This pattern may be due to a previously reported phenomenon in which low quality bases accumulate in certain genomic regions [18]; filtering reads by their minimum quality score may introduce a coverage bias. Regardless, MicrobeCensus includes options for filtering duplicate and low quality reads.

### Algorithm speed

Due to the increasingly large scale of metagenomics data, it is important to be able to process datasets in a reasonable time frame, particularly for users without access to high performance computing. For example, the Human Microbiome Project currently contains over 90 billion metagenomic reads spanning 1,298 samples [19]. Even a single lane on an Illumina HiSeq 2500 can generate over 500 Gb of sequence data [20].

Therefore, we benchmarked the speed of MicrobeCensus on a simulated 150-bp shotgun sequence library, which contained between 1,000 and 1 million reads, and compared these results to the speed of other methods run on the same data (Figure 2d). We found that MicrobeCensus was ∼320x faster than GAAS and ∼40x faster than our implementation of the Raes method. When using a single CPU, MicrobeCensus was able to process about 830 reads/second, or about 1 million reads in 20 minutes. Conversely, it took GAAS 10.2 hours to process only 100,000 reads. Using multi-threading we were able to further increase the speed of MicrobeCensus: 1.5x for 2 cores, 2.0x for 4 cores, and 2.5x for 8 cores. Most of the difference in speed between these methods is a result of the fact that MicrobeCensus searches reads against a database of only 30 genes, which together comprise less than 1% of most Bacterial genomes, and the difference in speed in the underlying sequence alignment algorithms – MicrobeCensus utilizes RAPsearch2, which is ∼20-90x faster than BLAST [21].

Additionally, we were interested in determining the fewest number of reads MicrobeCensus needed for a precise (i.e. low variance) estimate of AGS. For very large datasets, this would allow one estimate AGS using only a small fraction of total reads, thus greatly reducing the amount of required computational effort. On the other hand, this would enable one to exhibit caution in the case of smaller datasets, for which there may not be a sufficient number of reads for a precise, low variance estimate of AGS.

To address this question, we ran MicrobeCensus on several metagenomic samples from the Human Microbiome Project (HMP) [22] using between 10,000 and 20 million reads (see methods). For each sample, this allowed us to estimate the amount of dispersion (variance/mean) expected at a given number of sampled reads. For all samples, we found that dispersion was sufficiently low at 300,000 reads or more and reached an asymptote at about 500,000 reads, beyond which processing additional reads only marginally improved dispersion (Additional file 7). At sampling depths below 300,000 reads, dispersion quickly increased. Therefore, MicrobeCensus is able to generate low variance estimates of AGS from metagenomic datasets using as few as 300,000-500,000 reads while running in about 10 minutes on a single CPU.

### Average genome size is a source of bias for comparative analyses

We hypothesized that variation in AGS could significantly bias calculations of gene family relative abundance from shotgun metagenomes and impact downstream biological analyses. To address this question, we looked at the relative abundance of essential genes from the HMP and compared these values to estimates of AGS made from the same data. Specifically, we obtained standard estimates of relative abundance for KEGG Orthology Group (KOs) for 738 samples from the HMP Data Analysis and Coordination Center [23] and used MicrobeCensus to estimate the AGS of each corresponding metagenome (Additional file 8). We identified the KOs that corresponded to our set of 30 essential-single copy genes (Additional file 1) and compared the median estimated relative abundance of these genes to AGS across samples. As expected, we found a strong negative correlation between AGS and the relative abundance of essential single-copy genes among the stool communities (ρ = −0.84, p = 5e-37) (Figure 3). For example, the sample SRS013951 had an estimated AGS that was 0.48x that of SRS023346, and an estimated relative abundance of essential KOs that was 1.96x that of SRS023346. However, we did not find this relationship consistently among all body sites (Additional file 9). For example, we actually observed a slightly positive correlation within the oral communities (ρ = 0.29, p = 2e-8). This analysis prompted us to search for other sources of bias in relative abundance calculations.

**Figure 3.**
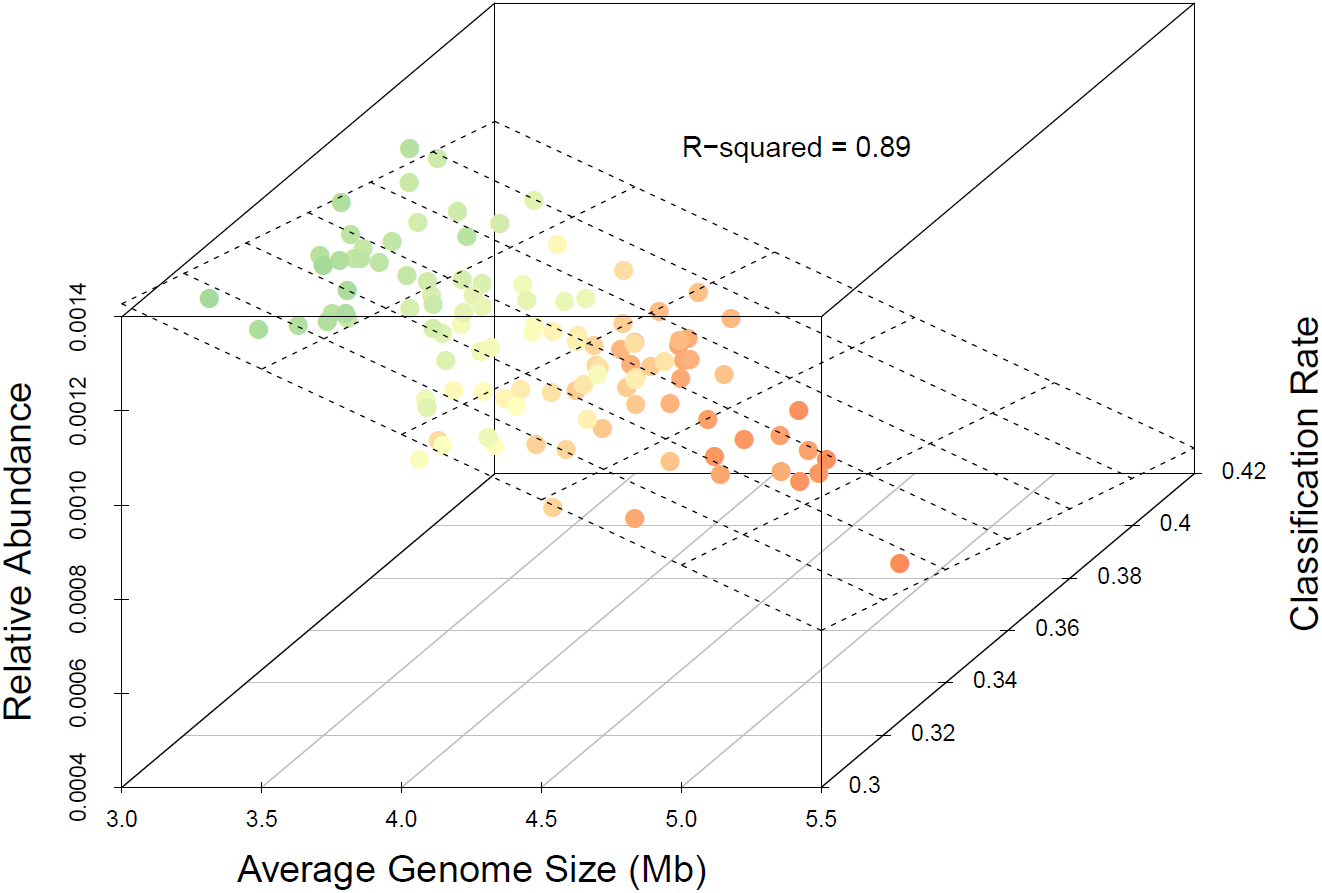
Average genome size is a source of bias for comparative analyses. The median relative abundance of essential, single-copy KOs versus AGS and classification rate for stool metagenomes from the HMP. Points are highlighted according to their relative abundance. Classification rate is defined as the fraction of reads classified into a KO. Fitting a plane to the data revealed that AGS and the classification rate are both anti-correlated with the relative abundance of essential genes and together explain 89% of the total variation.

We reasoned that, like a large AGS, a high classification rate (i.e. the proportion of the library that is assigned to any gene family) could artificially reduce relative abundance estimates. For example, essential single-copy genes could have a lower estimated relative abundance in a community with a higher classification rate, since these genes would make up a relatively smaller proportion of the total relative abundance of genes in that community. However, classification rates were not publicly available for HMP samples; to estimate this parameter, we searched 20,000 reads from each HMP sample against the KEGG database using BLASTX (see methods) (Additional file 10). Next, we corrected for AGS and then examined the relationship between classification rate and the median abundance of essential KOs in the HMP samples. As expected, we observed a strong negative correlation across each of the different body sites (−0.87 >= ρ >= −0.76) (Additional file 9). We also found that AGS and classification rate were highly anti-correlated (ρ = −0.71, p= 5e-103); communities with large AGS tended to have proportionally fewer reads classified into a KO, as one might expect if large genomes contain relatively more novel genes. Based on these findings, we re-examined the relationship between AGS and the median abundance of essential genes after correcting for differences in classification rate. This analysis revealed the expected negative correlation across all body sites, including stool (ρ = −0.94), oral (ρ = −0.72), nares (ρ = −0.79), urogenital tract (ρ = −0.90), and skin (ρ = −0.86) (Additional file 10). When considered together, AGS (p = 9e-63) and the classification rate (p = 1e-28) accounted for 89% of the total variation in essential KOs among the stool communities (Figure 3).

Thus, both classification rate and AGS can be major sources of bias when comparing the relative abundance of genes across samples and should be accounted for in addition to other known biases such as gene length and library size. Furthermore, we expect that this problem is not limited to the human microbiome, but to shotgun metagenomics studies in general. We suggest that normalizing (i.e. dividing) by the relative abundance of essential single-copy genes should eliminate these unwanted sources of variation and improve downstream biological analyses. We provide a list of the identifiers for these genes in the KEGG database (Additional file 1), and they are listed elsewhere for COGs [9]. It is also possible to use MicrobeCensus to directly estimate and normalize (i.e. multiply) for the AGS of a community – this is particularly important for comparative metagenomic studies that estimate the abundance of specific genes of interest or for databases where these 30 essential genes are not known.

### Average genome size varies systematically in human microbiome data

To further explore natural variation of genome size in the human microbiome, we used MicrobeCensus to estimate the AGS of over 1,300 metagenomic samples from human subjects, collectively spanning five major body sites (GI tract, skin, airways, mouth, and urogenital tract) and five countries (US, China, Sweden, Denmark and Spain) (Additional file 8). Included in this analysis were samples from the HMP and three other large studies of the gut microbiome [1, 5, 6] (see methods).

We found significant differences in AGS between nearly all body sites within the HMP data (Figure 4). The stool communities had an especially large and broad range of AGS (2.77 to 5.77 Mb, mean = 3.90 Mb). This finding is consistent with previous reports that stool communities have the highest ratio of genes per OTU [22], and that several prominent members of the gut have large genomes including: *B. thetaiotamicron* (6.5 Mb), *B. ovatus* (6.5 Mb), and *B. vulgatus* (4.9 Mb). The skin and nares communities had intermediate AGS, but each contained several outliers greater than 6 Mb – possibly due to a high relative abundance of microbial eukaryotes in these samples that has been previously reported [24]. The oral and urogenital communities were characterized by the lowest AGS (mean = 2.23 and 2.11 Mb, respectively), consistent with a high relative abundance of Streptococcus and Lactobacillus in these respective sites. For example, only 3 of the 396 oral samples had an estimated AGS that exceeded any of the 146 stool communities. Within the oral cavity, samples from the subgingival and supragingival plaques tended to have slightly larger AGS (mean = 2.5 to 2.6 Mb) compared to the tongue, throat, and buccal mucosa (all means < 2.2 Mb). Surprisingly, we found that AGS was remarkably stable within each of the oral sites; the maximum coefficient of variation in any of the oral sites was only 0.08, in contrast to high values in stool (0.16), urogenital tract (0.23), airways (0.36), and skin (0.45). The stability of AGS in the mouth may be due to a combination of factors, including lower beta-diversity and greater functional convergence than other body sites, although this needs to be investigated in greater detail. Interestingly, we also observed relatively large AGS variation in the urogenital tract, despite the fact that this community is dominated by various *Lactobacillus* species (spp). Comparing the distribution of stool AGS estimates across studies, we found significant differences between countries of origin (Sweden mean = 3.03 Mb, Spain mean = 3.17 Mb, Denmark mean = 3.27 Mb, China mean = 3.47 Mb, US mean = 3.95 Mb), which may reflect differences in age, diet, antibiotic use, or other environmental and ecological forces between the distinct continents and populations surveyed (Additional file 11).

**Figure 4.**
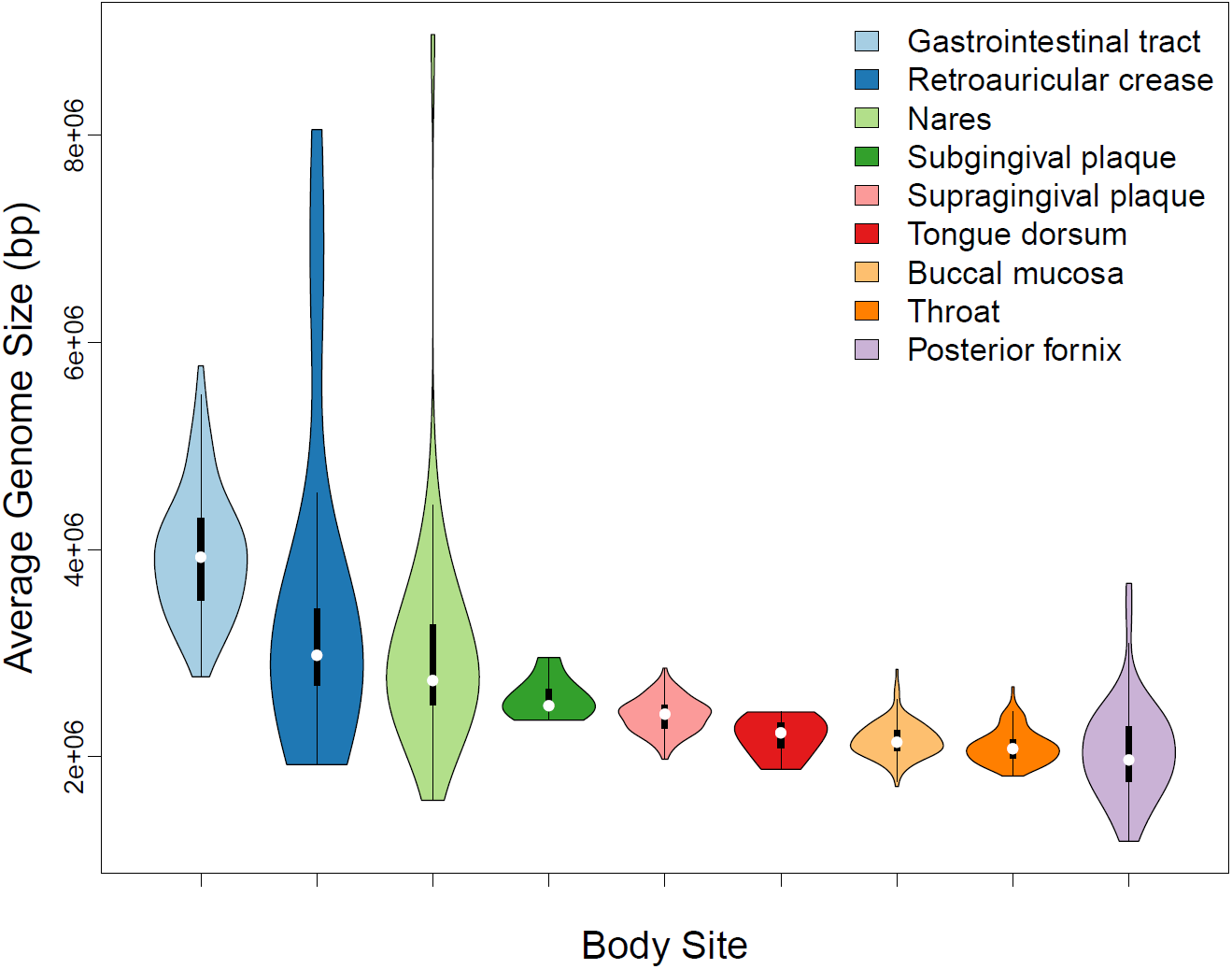
Average genome size varies systematically in human microbiome data. Distribution of estimated AGS for 736 samples from the Human Microbiome Project.

Perhaps unsurprisingly, we found that differences in AGS within and between communities could be largely explained by genus and species level taxonomic variation of Bacteria. Using a multiple-linear model of species-level relative abundance, we were able to explain 74% of the total variation in AGS across all HMP samples. Within the stool communities, we found that AGS was positively correlated with abundant *Bacteroides* spp. including *B. ovatus* (6.5 Mb), *B. thetaiotaomicron* (6.3 Mb), and *B. xylanisolvens* (6.0 Mb) (all p < 1e-4, all r^2^ > 0.12). However, most of the variation was explained by unclassified *Bacteroides* spp. (p = 6e-11, r^2^ = 0.36), potentially indicating the presence of other large *Bacteroides* genomes that have yet to be discovered in the gut microbiome. In contrast, members of the order Clostridiales (p = 1.5e-7, r^2^ = 0.18) were negatively correlated with genome size, consistent with reports that Firmicutes possess smaller genomes and a disproportionately smaller number of glycan-degrading enzymes than *Bacteroides* [25]. In the skin and nares communities, we found few strong taxonomic associations with genome size, and our multiple-linear model of Bacterial species explained relatively little AGS variation within these body sites (0.13 and 0.03 cross-validation r^2^, respectively). This suggests that the majority of genome size variation in these communities may not be driven by the relative abundance of Bacteria, but instead may be influenced by the abundance of fungi or viruses. Consistent with this hypothesis, we found that the relative abundance of microbial eukaryotes was as high as 30% in these communities. In the oral body sites, we found members of the genera *Mycobacterium* (r^2^ = 0.34), *Bifidobacterium* (r^2^ = 0.31), and *Actinomyces* (r^2^ = 0.33) were all positively correlated with AGS, while members of *Streptococcus* (*S. infantis* and *S. mitis*) and of *Haemophilus* (*H. influenzae* and *H. parainfluenza*) were negatively correlated with genome size. Lastly, while the posterior fornix was dominated by different *Lactobacillus* spp. (mean relative abundance = 0.91), variation in AGS was best explained by members of other genera, including: *Bacteroides*, *Parabacteroides*, *Alistpes*, and *Eubacterium*.

Interestingly, after accounting for taxonomic differences between samples, there were no longer any significant differences in AGS between body sites, and adding body site to the linear model of species abundance did not increase the amount of explained AGS variation. On the other hand, body site alone was able to explain 63% of total variation in AGS. This suggests that the differences in genome size between body sites are largely driven by differences in species abundance distributions, rather than differences in genome size between closely related taxa occupying different sites. For example, AGS is larger in stool communities than in oral communities mainly because stool communities have a greater relative abundance of *Bacteroides* (r^2^=0.83, p=0). This is consistent with previous observations that environmental filtering is a major ecological force in the human microbiome [8]. However, this finding does not exclude the possibility that there are also site-specific evolutionary forces that act to expand or contract genomes of the different taxa that colonize each site. It is likely that both of these forces – environmental filtering and evolution – have lead to the observed differences in genome size across the human microbiome. These and other possibilities need to be investigated in greater detail in future work.

### Average genome size variation reflects diverse modes of functional adaptation in the gut microbiome

Given that the distribution of AGS in the gut microbiome was so broad (2.77 to 5.77 Mb), we were interested in identifying the genes and pathways that were contributing to this large difference, particularly since genome size and gene count are directly proportional in Bacteria [13]. Based on genome analysis of sequenced gut isolates, it has been suggested that specialists tend to have smaller genomes than generalists and that organisms adapted to live outside the host have large accessory genomes [8]. Supporting these views, analysis of large *Bacteroides* genomes have revealed expanded repertoires of polysaccharide utilization genes that confer *Bacteroides* spp. metabolic flexibility and the ability to harvest both diet and host-derived glycans [26]. While these are interesting observations, to the best of our knowledge, the relationship between genome size and functional ecology has not been directly addressed in metagenomes from the human gut microbiome.

To shed light on the relationship between genome size and functional ecology in the gut, we searched for functions (genes, modules, and pathways) that were strongly correlated with AGS across stool communities within the HMP. Importantly, we first normalized the relative abundance of functions by the 30 essential KOs (Additional file 1) in order to eliminate bias from AGS and other sources. Specifically, for each sample we divided the relative abundance each KO by the median relative abundance of the 30 essential single-copy KOs for that sample. After normalization, we found that AGS tracked with major functional differences across stool samples. A principle component analysis of normalized KO relative abundance revealed that AGS was strongly correlated with the first principle component (r^2^ = 0.56, p < 1e-23) (Additional file 12). Furthermore, we found over that 1,700 KOs, 200 KEGG modules, and 150 KEGG pathways were strongly associated with AGS across stool samples (all p < 1e-10) (Additional files 13-14). Most of these functions tended to be more abundant in communities with larger genomes, as one might expect if larger genomes have a greater number of genes. For example, 78% of significant KOs were positively correlated with AGS. Furthermore, 45% of KOs associated with AGS in the HMP were independently replicated in large gut microbiome project from Chinese individuals [5], suggesting that the gene expansions we observe are not limited to the population studied by the HMP.

We found that our normalization procedure was critical to reveal both the magnitude and direction of functional variation among stool samples. Normalizing by essential single-copy KOs enabled us to identify twice the number of positively correlated genes (p < 1e-10, ρ>0) and prevented us from identifying 878 potential false positives, including each of the 30 essential single-copy KOs (all p = 0, mean ρ = −0.84). Many of the top-ranked pathways would have been missed using standard relative abundance to quantify gene families (Additional file 15). In fact, without proper normalization, the major direction of functional variation would have been reversed, with 68% of differentially abundant genes (p < 1e-10) instead being negatively correlated with AGS.

A striking pattern emerged when looking at the top-ranked pathways and modules that were either strongly positively or negatively correlated with AGS across stool communities – we found that positively correlated functions (i.e. more abundant in communities with large AGS) were dominated by metabolism, biosynthesis, and two-component signaling, while negatively correlated functions (i.e. more abundant in communities with small AGS) were dominated by various transport modules (Figure 5a-b). This finding is in agreement with a comparative genomics analysis of *Bacteroides* and *Firmicutes*, which found that *Bacteroides* possess larger genomes and a disproportionately greater number of glycan-degrading enzymes than *Firmicutes* [25]. Furthermore, this study found that the *E. rectale*, a prominent *Firmicutes* species in the gut, adapts to *B. thetaiotamicron* by decreasing production of its glycan-degrading enzymes, and increasing expression of selected amino acid and sugar transporters [25]. Together, these findings highlight both metabolic interactions and functional specialization in the gut microbiome. At the KO level, we found that, rpoE (K03088), an ECF-sigma transcription factor involved in regulating expression of polysaccharide utilization loci [26], was strongly correlated with AGS in both the HMP (ρ = 0.87) and Chinese microbiome project (ρ = 0.82) (Figure 5c). Additionally, rpoE reached extremely high levels of abundance in the gut (Figure 5c). Interestingly, deletion of this transcription factor in *B. thetaiotamicron* has been shown to produce defects in bacterial persistence and vertical transmission [26], further highlighting the importance of this gene in adaptation to the gut environment.

**Figure 5.**
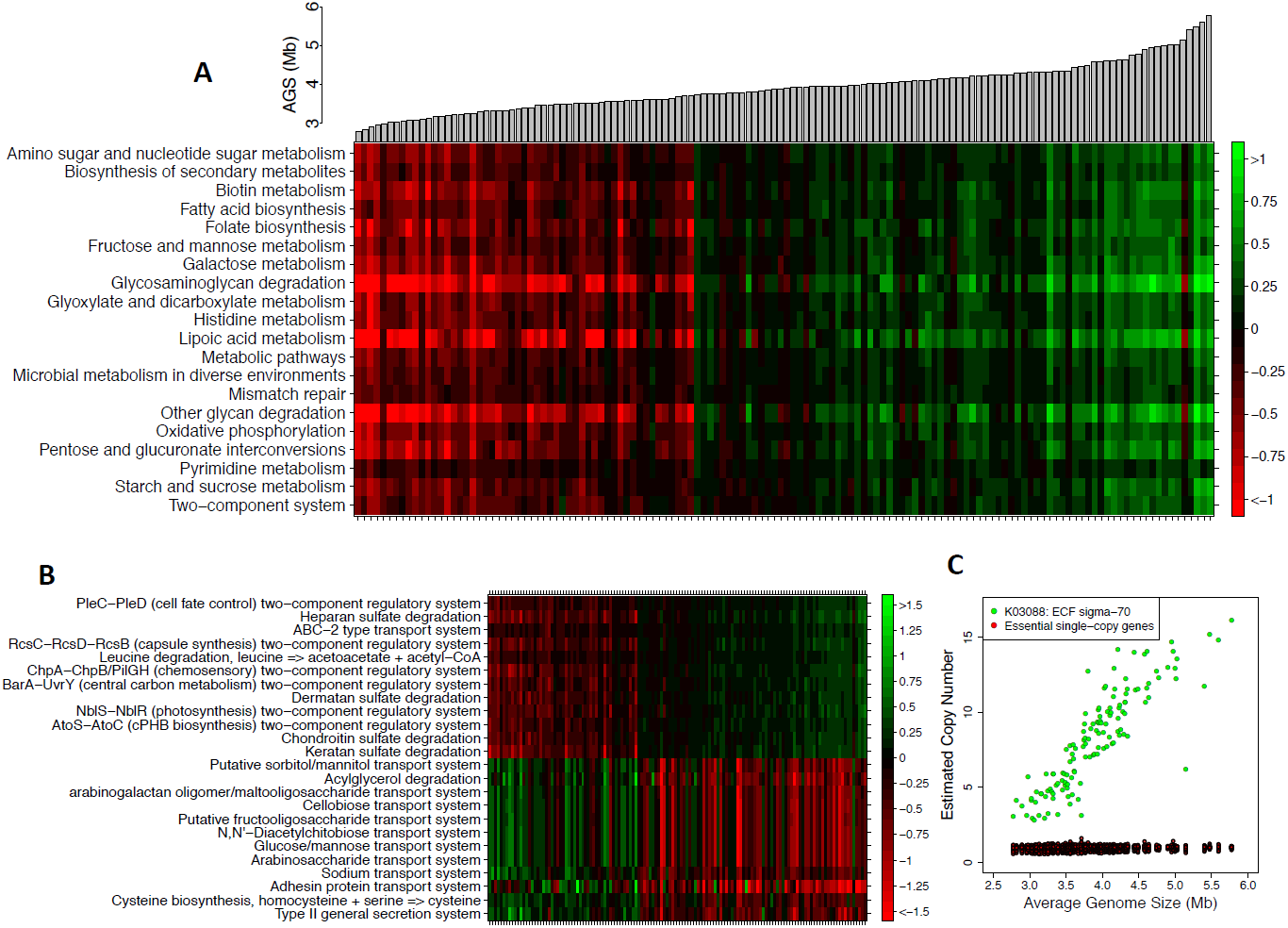
Average genome size reflects diverse modes of functional adaptation. Normalized relative abundance of functions (KOs, modules, and pathways) that track with AGS in the gut microbiome. **(a)** Log2 fold change of the top-20 KEGG pathways positively correlated with AGS across stool samples from the HMP. AGS is plotted above each sample at the top of the figure. **(b)** Log2 fold change of the top modules that are either positively or negatively correlated with AGS across stool samples from the HMP. **(c)** Normalized relative abundance of the ECF sigma-factor, K03088, and essential single-copy KOs across stool samples from the HMP.

## Conclusions

MicrobeCensus is a novel computational method that rapidly and accurately estimates the AGS of a microbial community from metagenomic data by aligning reads to a set of 30 essential single-copy genes. The accuracy of our method is largely due to extensive optimization of read length specific alignment parameters, which limit the technical variation of these genes across metagenomes. We performed careful validation of our method, demonstrating that unlike existing methods, MicrobeCensus is able to accurately estimate AGS for libraries of varying read length and for communities of varying phylogenetic divergence from sequenced organisms.

Applying MicrobeCensus to a large number of human microbiome samples, we confirmed that variation in AGS can significantly bias estimation of gene family relative abundance from shotgun metagenomes. Additionally, we found that the classification rate (i.e. the fraction of reads from a library that are classified into a protein family database) is also a major source of bias. Together, AGS and the classification rate accounted for 89% of variation of essential single-copy genes among stool samples from the Human Microbiome Project, which varied in relative abundance from 6 × 10^-4^ to 1.4 × 10^-3^.

Finally, we found that AGS varies systematically across body sites and individuals, and tracks with important functional and taxonomic differences in the microbiome. For example, in the gut, AGS ranges from 2.5 to 5.8 megabases and is positively correlated with the abundance of *Bacteroides*. After normalizing for AGS and other sources of bias, we found that communities with large AGS were enriched for pathways related to metabolism, biosynthesis, and two-component signaling, whereas communities with small AGS were enriched for functions related to transport of saccharides. These novel observations would have been missed without appropriate normalization.

We expect our approach to be widely applicable. MicrobeCensus is easy to use and can be applied to shotgun metagenomics data from any environment; it is not limited to studies of the human microbiome. We expect the improved accuracy of MicrobeCensus over existing AGS methods to be even greater in environments (e.g., soil, marine) where fewer genomes have been sequenced. In this study, we focused on shotgun sequenced DNA. In shotgun transcriptomics, similar normalization issues will arise, in conjunction with RNA-specific issues such as the large dynamic range of expression values. Since we expect the essential genes used by MicrobeCensus to be universally expressed, MicrobeCensus could easily be modified to adjust for and investigate AGS in metatranscriptome studies. To facilitate such extensions, our open-source software is freely available. This work comes at a critical time for the metagenomics field, as more and more studies attempt to quantify and compare relative amounts of many gene families and pathways across samples.

## Methods

### Essential, single-copy gene families

We downloaded a recently published set of 40 phylogenetically diverse, single-copy protein families found in nearly all Bacteria and Archaea [17]. To validate that these genes were in fact universally distributed and single copy, we searched these sequences against the proteomes of 250 Bacteria and 79 Archaeal genomes (Additional files 1-2) using BLAST [27]. An essential gene was deemed present in a genome if it’s corresponding E-value was below 1E-5 and if the query and target proteins were both covered by at least 70% of their length. We identified a subset of these families (N = 30) with a mean copy-number very close to 1.0 and copy-number variance close to zero. Nearly all of these genes (27/30) were components of the ribosome.

### Method training and optimization

Now that we identified a set of essential single-copy genes, we were interested in using these genes to estimate AGS from metagenomes. While we expect that the fraction of metagenomic reads that are assigned to an essential gene should be inversely proportional to AGS, this relationship depends on both read length and the similarity threshold used to identify homologs (Figure 6a). As shown in figure 6a, more reads are classified into an essential gene per genome equivalent in the short-read libraries relative to the long read libraries. On the other hand, fewer homologs will be detected at a strict sequence similarity threshold (e.g. bit-score > 60) relative to a more permissive one (e.g. bit-score > 30) (Figure 6a). Additionally, optimized, read-length specific similarity thresholds are required to minimize the technical variation of essential genes from metagenomic data (Figure 6b-c).

**Figure 6.**
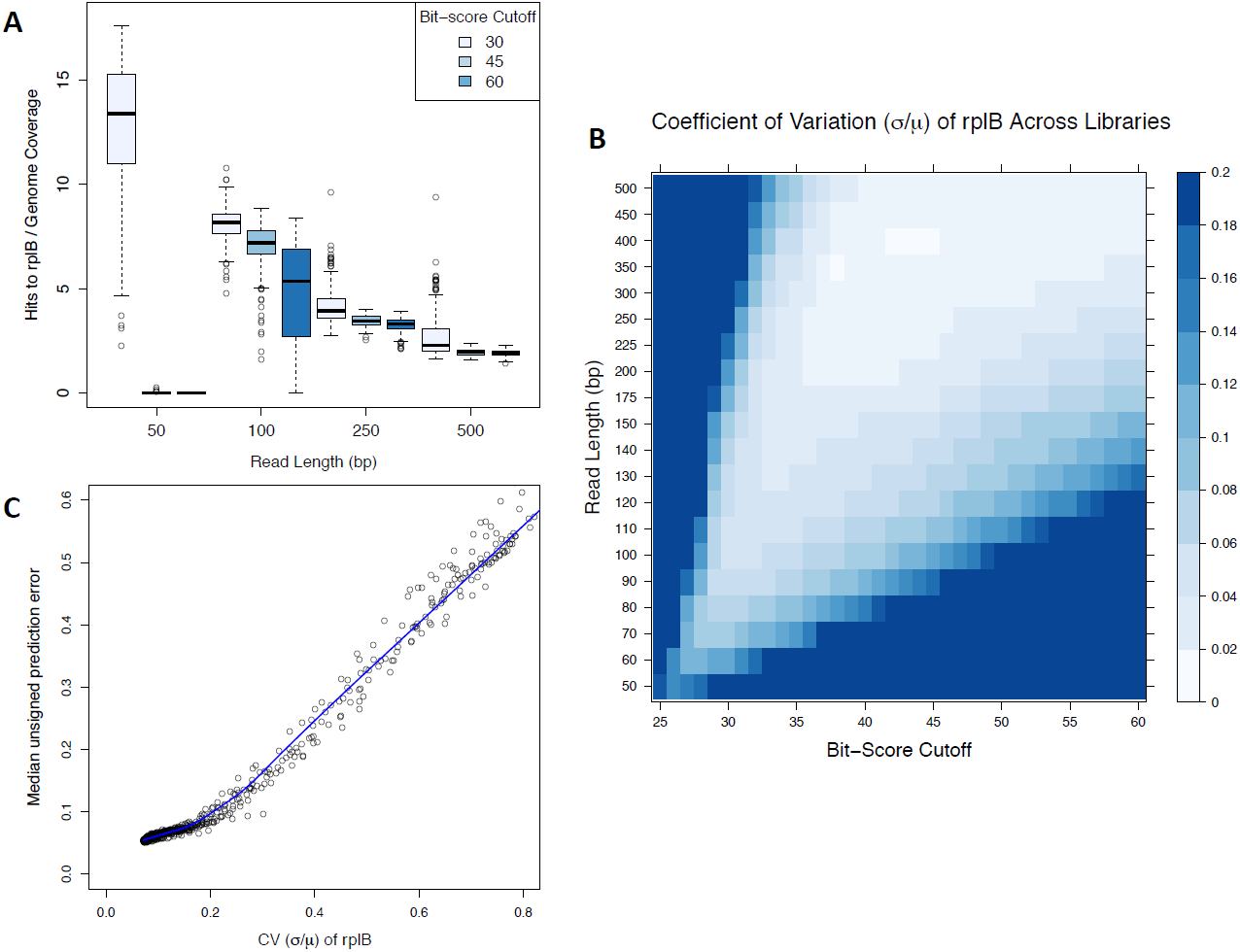
Optimal cutoffs minimize technical variation of essential genes across shotgun libraries. Reads from simulated shotgun sequence libraries were searched against the database of essential genes and classified at different cutoffs. Shown are results for one representative example, the essential gene – rplB. **(a)** The number of hits to rplB per genome equivalent (i.e. coverage) depends on both read length and the bit-score cutoff. Fewer homologs of rplB are detected in the short-read libraries (≤ 100 bp reads) and at more stringent cutoffs (≥ 60 bits). **(b)** The coefficient of variation of rplB (σ/μ) across simulated libraries depends on the bit-score cutoff. Read-length specific cutoffs are required to minimize the technical variation of rplB across all read lengths. **(c)** Picking cutoffs that minimize the variation of rplB also minimize the median unsigned error across simulated libraries when using rplB to estimate AGS.

Because these relationships cannot be determined *a priori*, we performed extensive shotgun sequencing simulations where genome size was known. Our goals were to identify optimal, read length-specific classification parameters, and to identify the expected relationship between genome size and the assignment rate at these cutoffs.

Toward this goal, we simulated shotgun sequence libraries from 329 complete Bacterial and Archaeal genomes (Additional file 2). We selected 250 Bacterial genomes from a large 16S gene phylogeny using an algorithm that maximized the total branch length of these taxa; additionally 79 Archaeal genomes were selected, each from a distinct genus. These genomes ranged in size from 138 Kb to 10 Mb and spanned 33 distinct phyla. All reference genomes were downloaded from the IMG database [28]. We used the software tool Grinder [29] to simulate shotgun sequence libraries from these genomes using read lengths ranging from 50 to 500 bp. Specifically, we simulated one 500-bp library for each of the 329 genomes at 200x coverage. For shorter read lengths, we simply trimmed the end off each read from these libraries to achieve the desired length. This resulted in 6,580 libraries and totaled over 400 Gb of sequence data.

Next, we searched each of these libraries against our database of essential genes using RAPsearch2 [21] (version 2.10; options: -z 1 -e 1 -t n -p f -b 0). A read was classified into an essential gene (at most one gene) if its alignment exceeded a set of specified criteria including: minimum bit-score, minimum alignment coverage, and maximum percent identity. We tested over 3,000 combinations of these cutoffs. At each read length, we identified the combination of cutoffs that minimized the technical variation of each gene across all 329 libraries (Figure 6b). Again, we expect these genes to be stable across libraries because each library contained the same number of genome equivalents (i.e. coverage), each gene was present in nearly all genomes, and each gene was present at only 1 copy per genome.

Finally, we used these optimized cutoffs to assign reads to the essential genes. This allowed us to identify the expected relationship between genome size and the fraction of a library assigned to each gene, *R*_*i,j*_, across the simulated libraries at each read length *i* and for each gene family *j*; this value is the proportionality constant, *Ĉ*_*i,j*_, in Figure 1. To assess performance on our training data, we estimated genome size for each simulated library: 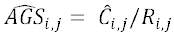. As expected, we found that the choice of classification parameters was critical for obtaining accurate estimates of genome size for our simulated libraries and that no set of parameters worked well for all read lengths (Figure 6b-c). The choice of a minimum bit-score threshold was particularly important. Relatively stringent bit-scores (>= 60 bits) worked well (< 1.8% median unsigned error) for reads longer than 300 bp, but failed miserably for reads shorter than 100 bp. Likewise, relatively lenient bit-scores (>= 30 bits) worked well (2.4–7.2% median unsigned error) for reads shorter than 100 bp, but were not accurate for longer reads. Overall, we were able to obtain accurate estimates of genome size for our simulated libraries (1.8–7.2% median unsigned error) for read lengths ranging from 50 to 500 bp.

### Comparison to existing methods

We compared the prediction accuracy of MicrobeCensus to GAAS (v0.17) for 20 simulated shotgun libraries, chosen at random from the set of 329 training genomes (Additional file 3). Each library was composed of 150-bp reads from a single organism and sequenced at 60x coverage.

For GAAS, we first searched the reads against the database of microbial genomes included with the software (NCBI RefSeq Release 56) using BLAST (v2.2.26) with the following options: blastall -m 8 -p blastn. We supplied the BLAST output to GAAS using the following command: gaas –f query.fna –d reference.fna –m blast.m8 –a taxon.map –v nucleic -e 1e-03 -p 0 -q 70 -sm 0 -j 1 -gp 0 -gs 0 -gt 0. We repeated this procedure at different levels of “taxonomic exclusion” in order to simulate libraries composed of organisms from novel taxonomic groups. Specifically, we discarded alignments from the BLAST output file where the target reference genome originated from the same taxonomic group (i.e. species, genus, etc.) as the organism in the simulated dataset.

In parallel, we ran MicrobeCensus using the same 20 simulated libraries at different levels of taxonomic exclusion. Specifically, we discarded alignments where the target essential gene originated from the same genome as the organism in the simulated dataset. Additionally, we retrained our linear models while excluding all training data that originated from the same taxonomic group as the organism in the simulated dataset. For both methods, we computed the prediction error as: |predicted size – expected size|/ expected size.

Separately, we evaluated whether the method described in Raes et al. was able to accurately estimate genome size for libraries composed of short-read data. To this end, we simulated shotgun libraries for the same 20 genomes used to benchmark GAAS. However, here, we simulated libraries with read lengths ranging from 50–500 bp, where each library was simulated at ≥ 20x coverage.

However, because no software was released, we had to manually implement the method described by Raes et al. Therefore, we downloaded version 3 of eggNOG database [30] and identified all protein sequences that corresponded to the 35 single-copy orthologous groups (OGs) used by Raes et al. Next, we created a BLAST database of these sequences using the tool formatdb and searched each simulated library against the 35 OGs with BLASTX (v2.2.26). We only assigned a shotgun sequence to one of the 35 OGs if the alignment score exceeded 60 bits but the percent identity was less than or equal to 50%. Finally, we computed marker density (x) as the number of hits per Mb of input data, and used the Raes formula to estimate genome size: Genome size = a+(b*L^-c)/x, where a = 21.2, b = 4230, c = 0.733, and L = library read length. We ran MicrobeCensus on the same data using default parameters and computed prediction error as above.

### Prediction accuracy using real data

To evaluate the ability of MicrobeCensus to estimate genome size from libraries composed of real data, we downloaded short-read Illumina datasets from 42 completed microbial genome projects using SRAdb [31] (Additional file 4). We specifically chose projects where the sequenced organism was not used to train our method. After converting the SRA files to FASTQ format, we used FASTQC [32] to identify adapter contamination in each dataset, and used cutadapt [33] (-e 0.1 --discard -O 5) to remove any contaminated sequences. Additionally, we removed exact duplicates using the first end of each paired-end read, and filtered reads with an average quality score of less than 5.

Next, we pooled this data to create ten mock metagenomes (Additional file 5). Each metagenome was composed of 5 million 70-bp reads from ten randomly selected genome projects (i.e. 500,000 reads from each project). We used MicrobeCensus to estimate AGS for each of these ten mock metagenomes. Finally, we evaluated the effect of various quality control procedures on AGS prediction accuracy for the 42 isolate genome projects. Specifically, we evaluated the effect of adaptor filtering, duplicate filtering, filtering by mean read quality, filtering by minimum read quality, and filtering reads with ambiguous base-calls (i.e. Ns) (Additional file 6).

### Speed benchmarking

We benchmarked the speed of MicrobeCensus, GAAS, and the Raes method on a simulated 150-bp shotgun sequence library, which contained between 1,000 and 1 million reads. GAAS was run with default options, except -v nucleic -sm 0, to increase its speed. MicrobeCensus was run with default options, except –t to specify the number of threads to use. The Raes method was run as previously described. All tests were performed on a server with 16 Intel Xeon X5560 2.80 GHz CPUs and 200 Gb of RAM running Ubuntu 10.04.4 LTS.

### Minimum number of reads for precise AGS estimates

To determine the minimum number of reads needed to precisely estimate AGS, we ran MicrobeCensus using different numbers of randomly selected reads from several real datasets. Specifically, we chose Human Microbiome Project samples from eight different body sites (SRS011084, SRS017849, SRS023847, SRS019029, SRS042984, SRS043663, SRS019127, SRS023468). For each sample, we selected up to 20 million (M) unique, single-end reads trimmed to 70 bp and searched these reads against the database of essential genes using RAPsearch2. Next, we used bootstrapping to repeatedly estimate AGS with MicrobeCensus using random subsets of these 20M reads. We evaluated sample sizes of 10,000 to 20,000,000 reads. For each sample size we performed 100 bootstrap iterations. This enabled us to estimate the amount of dispersion (variance/mean) observed for each sample at a given number of sampled reads (Additional file 7).

### Human microbiome sequence data

We downloaded 738 metagenomic samples from the Human Microbiome Project on September 22, 2013 from the HMP Data Analysis and Coordination Center (HMPDACC) [23]. Additionally, we downloaded metagenomic data for Chinese individuals [5] from the NCBI Sequence Read Archive [SRA045646, SRA050230] on December 22, 2013; for Spanish and Danish individuals from the MetaHIT project [1] from the Beijing Genomics Institute [ftp://public.genomics.org.cn/BGI/gutmeta] on January 13, 2014; and for Swedish individuals [6] from the NCBI Sequence Read Archive [ERP002469] on February 7, 2014.

### AGS for human microbiome samples

We used MicrobeCensus to estimate AGS for human microbiome samples using up to 5e6 single-end reads trimmed to 70 bp (Additional file 9). Additionally, we used options to remove duplicates (-d), filter reads with mean quality less than 5 (-m 5), and filter reads containing more than 5% unknown base calls (-u 5). In all downstream analyses, we only considered samples for which there were at least 300,000 reads after our quality control procedures.

### Abundance of KEGG Orthology groups

We downloaded HUMAnN [34] functional annotations for Human Microbiome Project samples on December 21, 2013 from the (HMPDACC) [23]. For the Chinese samples, we downloaded functional annotations from GigaDb [35] on December 20, 2013. All functional annotations were in the form of per-sample relative abundances of KEGG Orthology groups (KOs). Unfortunately, classification rates (% of reads mapped to the database) were not publicly available for HMP samples. To estimate classification rate, we searched 20,000 randomly selected reads from each HMP sample against the KEGG Orthology database using BLASTX (version 2.2.28+) and classified a read into a KO if it’s alignment E-value was less than 1.0, which was the same cutoff used by HUMAnN.

### Abundance of taxonomic groups

We downloaded MetaPhlAn [36] Bacterial and Archaeal taxonomic annotations for Human Microbiome Project samples on December 21, 2013 from the HMP DACC [23]. For other stool microbiomes, we ran MetaPhlAn version 1.7.7 using the options: --bowtie2db bowtie2db/mpa -- input_type = multifastq -t rel_ab. Additionally, we downloaded domain level taxonomic annotations for HMP samples on December 21, 2013 from the HMP DACC.

#### List of Abbreviations

AGS: average genome size of a microbial community
CV: coefficient of variation
Mb: megabase
CPU: central processing unit
NCBI: National Center for Biotechnology Institute
HMP: Human Microbiome Project
T2D: type-2 diabetes metagenomics sequencing project
MetaHIT: Metagenomics of the Human Intestinal Tract
OTU: operational taxonomic unit
KEGG: Kyoto Encyclopedia of Genes and Genomes
KO: KEGG Orthology Group

## Authors Contributions

SN wrote the software and performed the experiments and analysis. SN and KSP designed the experiments and wrote the paper. Both authors read and approved the final manuscript.

## Additional files

Additional file 1 is a table listing the 30 essential single-copy genes used to estimate average genome size. Additional file 2 is a table listing the 329 phylogenetically diverse Bacterial and Archaeal genomes used in shotgun sequencing simulations. Additional file 3 is a list of the 20 randomly selected genomes used to compare MicrobeCensus to existing methods. Additional file 4 is a table listing the NCBI short-read genome projects used for method validation and metagenome construction. Additional file 5 is a table listing the community composition of the ten mock metagenomes constructed from the short-read genome projects. Additional file 6 is a figure that illustrates the effect of various sequence quality filters on AGS prediction accuracy. Additional file 7 is a figure that shows the effect of sequencing depth on dispersion of AGS estimates from real metagenomes. Additional file 8 is a table listing AGS estimates for human microbiome data. Additional file 9 is a figure that shows the relationship between AGS, the classification rate, and the relative abundance of essential single-copy genes across all HMP body sites. Additional file 10 is a table listing classification rates (i.e. the fraction of library assigned to any KO) for HMP samples. Additional file 11 is a figure that shows the distribution of AGS across healthy subjects from the US, China, Denmark, Spain, and Sweden. Additional file 12 is a figure that shows the relationship between AGS and the first three principle components from a principal components analysis of normalized KO abundance across HMP stool samples. Additional file 13 is a table listing correlation coefficients for functions associated with AGS across body sites within the HMP. Additional file 14 is a table listing modules and pathways that are enriched for KOs that are significantly correlated (either positively or negatively) with AGS within body sites. Additional file 15 is a figure that shows the effect of normalization on identifying the KEGG pathways that are most positively associated with AGS.

## Acknowledgements

We thank Jonathan Eisen and Thomas Sharpton for providing valuable discussion. We are grateful to Dongying Wu for providing a script we used to select a diverse set of taxa from a phylogenetic tree. This project was supported by funding from NSF grant #DMS-1069303, Gordon & Betty Moore Foundation grant #3300, the San Simeon Fund, and institutional funds from Gladstone Institutes.

